# Vertical distribution and light energy capture by AAP bacteria along contrasted areas of productivity in the Atlantic Ocean

**DOI:** 10.1101/2023.11.28.568994

**Authors:** Carlota R. Gazulla, Michal Koblížek, Jesús M. Mercado, Josep M. Gasol, Olga Sánchez, Isabel Ferrera

## Abstract

Aerobic anoxygenic phototrophic (AAP) bacteria are a common part of microbial communities in the sunlit ocean. They contain bacteriochlorophyll *a*-based photosystems that harvest solar energy for their metabolism. Across different oceanic areas and regimes, AAP bacteria seem to be more abundant in eutrophic areas, associated to high chlorophyll concentrations. While most of these studies are based on surface samples, little information is available on their vertical distribution in euphotic zones of the major ocean basins. We hypothesize that AAPs will follow a similar structure to the chlorophyll depth profile across areas with different degrees of stratification. To test this hypothesis, we enumerated AAP cells and determined bacteriochlorophyll *a* concentrations along the photic zone of a latitudinal transect in the South and Mid Atlantic Ocean. We show that the distribution of AAP bacteria is highly correlated to the chlorophyll *a* concentration and the abundance of picophytoplankton across vertical and horizontal gradients. Furthermore, we estimate the light energy captured by AAP bacteria across the water column and find that, while they share a common latitudinal pattern of light capture with the picophytoplankton, they display a unique vertical arrangement with highest photoheterotrophic activity is in the surface ocean.

## Introduction

The sunlit ocean, where there is enough light to sustain primary production, hosts a vast array of microorganisms that drive biogeochemical transformations and sustain life in the global ocean. Phytoplankton and other primary producers provide a continuous supply of dissolved organic matter that supports the growth of heterotrophic bacteria, that incorporate carbon into the base of the microbial food web (Azam et al. 1983). Besides the purely autotrophic and heterotrophic organisms, photoheterotrophic organisms are also key elements of the planktonic food chain since they are capable of harvesting light but rely heterotrophically on organic matter for growth. Alongside proteorhodopsin (PR)-containing bacteria, aerobic anoxygenic phototrophic (AAP) bacteria have attracted the interest of microbial ecologists since they were discovered to be widespread in the surface ocean (Kolber et al. 2000). AAP bacteria usually represent 1 to 7% of total bacteria in the euphotic zone (Koblížek 2015), but their larger size compared to other bacteria (Sieracki et al. 2006) and higher growth rates (Koblížek et al. 2007; Ferrera et al. 2011, Fecskeová et al., 2021) make them important components of the carbon cycle in the ocean.

AAP bacteria are close relatives of the purple non-sulfur bacteria as both contain bacteriochlorophyll-based reaction centres, although AAP bacteria have a purely aerobic lifestyle and cannot fix carbon. The photoheterotrophic metabolism of AAPs is based on the light harvesting pigment bacteriochlorophyll-*a* (Bchl *a*) that facilitates synthesis of ATP, and on organic substrates for growth. This advantageous metabolic combination led to the suggestion that AAP bacteria would be prevalent in oligotrophic ocean regions (Kolber et al. 2000). However, subsequent studies showed that AAP bacteria are actually more abundant in productive regions or shelf seas (Cottrell et al. 2006; Mašín et al. 2006; Sieracki et al. 2006; Hojerová et al. 2011; Ritchie and Johnson 2012, Vrdoljak et al., 2019), coastal lagoons (Lamy et al. 2011), and estuaries (Schwalbach and Fuhrman 2005; Cottrell et al. 2010), where they can account for up to 12% of the total prokaryotic community. In contrast, in oligotrophic sites such as the Sargasso Sea or the North Pacific gyre their relative contribution is low (<2%, Cottrell et al. 2006; Sieracki et al. 2006; Jiao et al. 2007; Gazulla et al. 2022). While AAPs are present throughout the entire epipelagic zone, most studies have focused on surface samples. The few studies that have dealt with their vertical distribution in the ocean showed that AAP bacteria had abundance trends different than those observed for heterotrophic bacteria or cyanobacteria (Cottrell et al. 2006; Sieracki et al. 2006; Jiao et al. 2007; Lami et al. 2007). Across the Mediterranean Sea, higher AAP abundances (Hojerová et al. 2011) and Bchl *a* concentrations (Gómez-Consarnau et al. 2019) were observed above and overlapping the deep chlorophyll maxima (DCM). Altogether, these results suggest that AAP bacteria could be associated to phytoplankton not only in the ocean surface but also along its water column.

Additionally, and in contrast to the growing knowledge on the distribution of AAP bacteria, there is little information regarding their phototrophic activity (i.e. light capture) in the natural environment. Laboratory studies with AAP cultures showed that light exposure reduced their respiration rate (Koblížek et al. 2010; Hauruseu and Koblízek 2012), and increased the efficiency of carbon metabolism (Piwosz et al. 2018). In result, the cultures grown on light and dark regime accumulated more biomass than cultures kept in the dark (Hauruseu and Koblížek, 2012, Piwosz et al. 2018). Light stimulation of growth of natural populations of AAP bacteria was also demonstrated in field experiments (Ferrera et al. 2017, Sánchez et al. 2020). In addition, theoretical estimates indicate that AAP bacteria harvest more light energy per cell than PR-containing bacteria (Kirchman and Hanson 2013). However, estimates of actual energy fluxes related to light in the marine environment are scarce. By determining AAP abundance and pigment concentration we can determine the light energy captured by AAP cells, providing information on their *in situ* physiology and, ultimately, allowing the estimation of their potential phototrophic activity.

We studied in detail the distribution of AAP bacteria along the water column in areas of contrasting productivity across the South and Mid Atlantic Ocean. We took samples in 27 stations at different depths following the fluorescence variation depicted by the CTD profiler and enumerated the abundance of AAP bacteria, as well as that of other photosynthetic microorganisms. We hypothesize that the distribution of AAP bacteria along the water column will be strongly connected to the DCM structure, and with higher abundances at more productive sites. In addition, we used an improved high-performance liquid chromatography (HPLC) protocol to accurately detect picomolar (pM) concentrations of Bchl *a* in the water column. Based on the microscopic and pigment data, we estimated the light energy captured by both Bchl *a* and Chl *a* and described how this varied spatially and along the depth gradient.

## Material and Methods

### Sampling

The POSEIDON Expedition took place between March and April 2019, on board the Spanish *RV Sarmiento de Gamboa*. It covered a distance of ∼9,000 km across a latitudinal transect (48°S/26°N) in the Atlantic Ocean. A total of 27 stations were sampled at different depths along the epipelagic zone. A SeaBird 911-plus CTD profiler was used to profile temperature, salinity, conductivity, fluorescence, dissolved oxygen, and other variables. The fluorescence values from the CTD allowed us to define the DCM structure along the water column and, according to this structure, in each station we took samples at the surface, above the DCM, at the DCM peak, below the DCM and at the shallowest depth where chlorophyll fluorescence was undetectable. Six of these stations were sampled at a higher resolution (10 depths across the DCM structure). At two stations the DCM structure displayed two peaks of chlorophyll maxima and samples were taken at each peak as well as between peaks (stations 10 and 16). Stations are numbered from 1 to 28, yet there are no samples from station 15 in our dataset. In total, we collected 172 samples from different depths ranging from 5 m to 280 m. All seawater samples were prefiltered by a 200-μm mesh to remove large plankton.

### Cell abundances

Samples (1.6 mL) were fixed using a solution of 1% paraformaldehyde + 0.05% glutaraldehyde (final concentrations), deep frozen in liquid nitrogen and stored at −80 °C until analysed. Prokaryotic and photosynthetic picoeukaryotic abundances were determined by flow cytometry as described in Gasol and Morán (2015). Additionally, total bacteria (i.e. prokaryotes) and AAP bacteria were counted using infra-red epifluorescence microscopy (Mašín et al. 2006) with recent modifications (Piwosz et al. 2022). Nine mL of seawater were fixed with formaldehyde to a final concentration of 3.7% and filtered onto white 25 mm polycarbonate filters of 0.2 µm pore size (Nuclepore; Whatman). Cells were stained with 4’6-diamidino-2-phenylindole (DAPI) and counted using an epifluorescence Zeiss Axio Imager.D2 microscope equipped with a Plan-Apochromat 63×/1.46 Oil Corr objective and a Collibri LED module illumination system (Carl Zeiss, Jena, Germany). Images for DAPI fluorescence (total bacteria) were taken under blue emission, the autofluorescence of Chl *a* was recorded in the red part of the spectrum and both AAP bacteria and Chl *a*-containing bacteria were recorded in the infrared part of the spectrum. To obtain net AAP bacterial counts, the contribution of Chl *a*-containing organisms to the infrared image was subtracted (Cottrell et al. 2006). We took at least 10 microphotographs for every sample and analysed them with the ACMEtool2.0 software (Bennke et al. 2016).

### Pigment analyses

Pigment composition in seawater samples was analysed using an improved high-performance liquid chromatography (HPLC) protocol derived from the method of Goericke and Repeta (1993). Between 1 and 2 L of seawater were prefiltered through a 200 μm mesh, and then collected onto 25 mm polyethersulfone (PES) 0.45 μm pore size filter discs (Meck Millipore Ltd.) to assure that all AAP cells were captured (Lami et al. 2009). The use of 25 mm PES filters also reduced the volume of filtered seawater, reduced the amount of extraction solvents needed, and simplified the extraction protocol. Filters were conserved at −80°C until pigment extraction which was conducted in the laboratory. Filters were transferred to cryogenic vials with 1 mL of 7:2 acetone:methanol (v/v) mixture and ∼30 mg of glass beads, and then disintegrated in a Mini-Beadbeater™ (Biospec Products, USA) during 2 minutes, to break the filters and extract the pigments. After a short centrifugation (3 min, 14.100 rcf), the clear extracts were transferred to HPLC vials for further processing. We used a SCL-40 HPLC (Nexera series, Shimadzu, Japan) equipped with SPD-M40 PDA detector and a UV-VIS detector with a deuterium (D2) lamp. For each sample, 50 μL were injected into the system. Pigments were separated using a heated (40°C) Kinetex® 2.6μm C8 100 Å column (Phenomenex Inc., CA, USA) with binary solvent system consisting of A: 75% methanol + 25% 28 mM ammonium acetate, and B: 100% methanol, with a flow rate of 1 mL·min^‒1^. The gradient was the following: A/B 100/0 (0min), A/B 0/100 (21 min), 0/100 (23 min), 100/0 (24 min), and 100/0 (25 min). The peak for BChl *a* was registered at 770 nm. The pigment concentration in the original samples was calculated from the peak area. The use of Kinetex® column with core shell technology resulted in sharper peaks, which helped to accurately integrate small BChl *a* peaks. The peak of chlorophyll *a* and divinyl-chlorophyll *a* were integrated together at 656 nm, so values of Chl *a* presented in the results include both chlorophyll *a* and divinyl-chlorophyll *a*. The HPLC system was calibrated using 100% methanol extracts of *Synechocystis* sp. PCC6803 for Chl *a* and *Rhodobacter sphaeroides* for BChl *a*, where the concentrations of Chl *a* and Bchl *a* were determined spectroscopically using a Shimadzu UV2600 spectrometer. From the Bchl *a* concentration and AAP cell abundances, we calculated the Bchl *a* cell quota as the quantity of Bchl *a* per cell.

### Satellite-derived variables and irradiance per depth calculations

Given that no in situ PAR data was measured during the cruise, satellite-derived photosynthetic active radiation (PAR) and diffuse attenuation coefficients at 490 nm (Kd) were obtained from the Suomi-NPP/Visible Infrared Imaging Radiometer Suite instrument (SNPP-VIIRS). Eight-day composites at 9.28 km resolution (level 3-mapped data) were downloaded from the NASA Ocean Color Web (https://oceancolor.gsfc.nasa.gov/l3/). Both variables for each station were extracted according to the sampling date and the station coordinates. In case of data gaps due to clouds, the average values of a 10×10 pixel square centred on the station coordinates were calculated. In situ irradiance per depth (*PAR_(Z)_*) was calculated as:

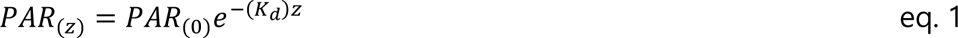

where *PAR_(0)_* is the subsurface average daily downward irradiance for each station, assuming an average surface reflectance of 6% (Bailey et al., 2008), *z* is the depth in meters and *K_d_*is the diffuse attenuation coefficient determined for each station. In equations 2 and 3 below, *PAR_(Z)_* is denoted as *I* (irradiance).

### Bioenergetic calculations

The cellular daily energy captured per each cell was estimated applying the theoretical calculations of Kirchman and Hanson (2013) for AAP bacteria, assuming a hyperbolic response to light, as it was also estimated in Gómez-Consarnau et al. (2019). First, we calculated the number of photosynthetic units (PSU) per cell, considering that AAP bacteria contain 34 molecules of Bchl *a* (Yurkov and Beatty 1998), and assuming an average number of 300 molecules of Chl *a* in the case of the autotrophic picoplankton as in Gómez-Consarnau et al. (2019). We used AAP abundances obtained through microscopic analyses, and the abundances of the autotrophic picoplankton estimated as the sum of the abundances of *Prochlorococcus*, *Synechococcus*, and picoeukaryotes from the flow cytometry analyses. These calculations were applied to stations 5 to 28, and we decided to remove stations 1, 2, 3 and 4 from the analysis, since we assumed that a large fraction of the phytoplankton in samples from those stations are diatoms and could alter our estimations. As suggested by Kirchman and Hanson (2013), we considered that electron fluxes vary with irradiance in a Michaelis-Menten type relationship as:

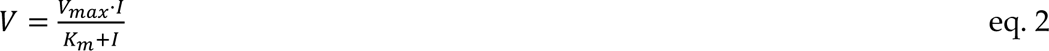

where V_max_ is the maximum flux and K_m_ is the half saturation constant, that is, the value of irradiance (*I*) at which V is half of V_max_. The values of V_max_ and K_m_ used in the equations, as well as the energy per photon for Bchl *a* and Chl *a* were extracted from Kirchman and Hanson (2013).

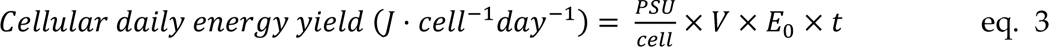

The cellular daily energy yield was estimated as in equation 3, where E_0_ is the energy per photon and *t* is 43200 seconds, as we considered 12 h of light per day.

All calculations were performed in the R Core v4.2.0 (R Core Team 2022) environment, using *tidyverse* (Wickham et al. 2019) and *ggplot2* (Wickham 2016) packages for data processing and figure representations. Besides, Ocean Data View v5.6.5 was used to represent sectional distribution images.

## Results

### 1. Oceanographic context

The transect went from south to north starting in a region of high productivity in the South Atlantic, characterized by low temperatures (mean 9.3 °C) and lower salinity values (mean 34.2), influenced by their proximity to the Southern Ocean (stations 1 and 2, beyond 40° S). Fluorescence and nutrient concentrations were high in these southernmost stations, especially silicate (mean 5.53 µM in this area vs. 2.15 µM in the whole transect). Noteworthy, in the HPLC analyses of samples from these stations we observed high values of fucoxanthin (data not shown), the characteristic pigment of diatoms. In subsequent stations going north, temperature and salinity increased up to the maximum values recorded in the transect (temp. >28.5 °C and salinity >37 in stations 10 to 14). These stations, located within the South Atlantic oligotrophic gyre (Figure 1), had very low values of surface fluorescence, and the DCMs appeared below 120 m depths (Figure 2A). The lowest values of nitrite, nitrate, phosphate, and silicate were recorded in the oligotrophic gyre (Figure S1). Interestingly, in this area, we observed two stations with a two-peak DCM structure (station 10 and station 16). In these cases, the shallowest peak was located at ∼40 m and ∼25 m depth respectively, and the deeper one, at depths of ∼100 and ∼80 m, had the highest values of fluorescence (Figure 2A). As the cruise progressed towards the equator, the DCMs were shallower (between 50-100 m deep) and characterized by higher values of fluorescence and nutrient concentrations, while temperature and salinity decreased. Finally, beyond the equator, seawater temperatures remained at around 20 °C but salinity increased as we were approaching the North-Atlantic oligotrophic gyre. Stations 23 to 28 had low nutrient levels and strong vertical mixing due to the upwelling waters off the coast of Mauritania, being the mixed layer depth (MLD) below 100 metres in several of them.

**Figure 1.**
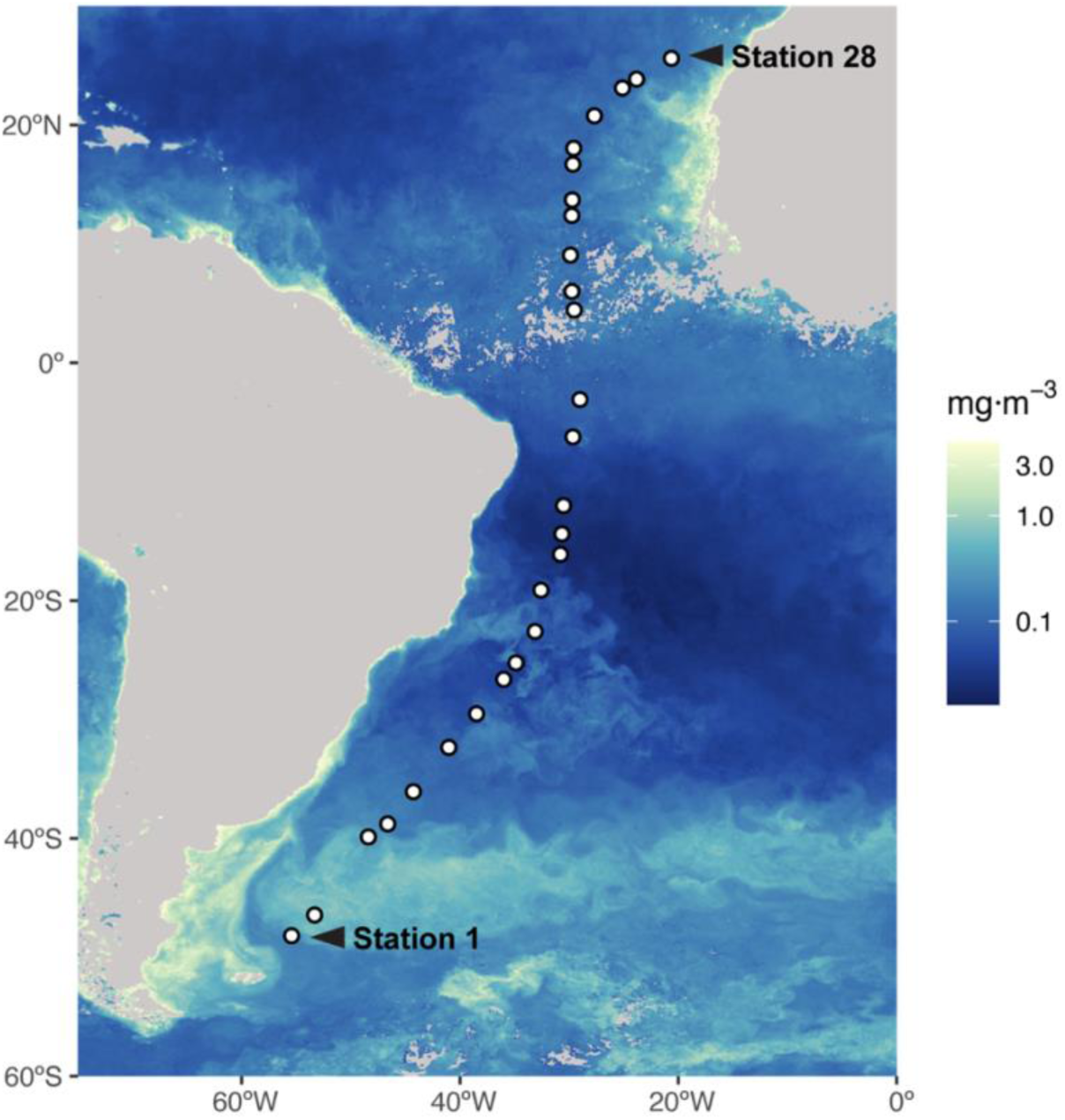
Near chlorophyll surface concentration. Transect of the Poseidon Expedition along the South and Mid Atlantic Ocean. Colours represent the near-surface chlorophyll concentration (mg·m^-3^) in March 2019. Data extracted from https://oceancolor.gsfc.nasa.gov/l3/ from the SNPP-VIIRS instrument. Each dot represent one station, numbered from 1 to 28. Note that station 15 does not exist.

**Figure 2.**
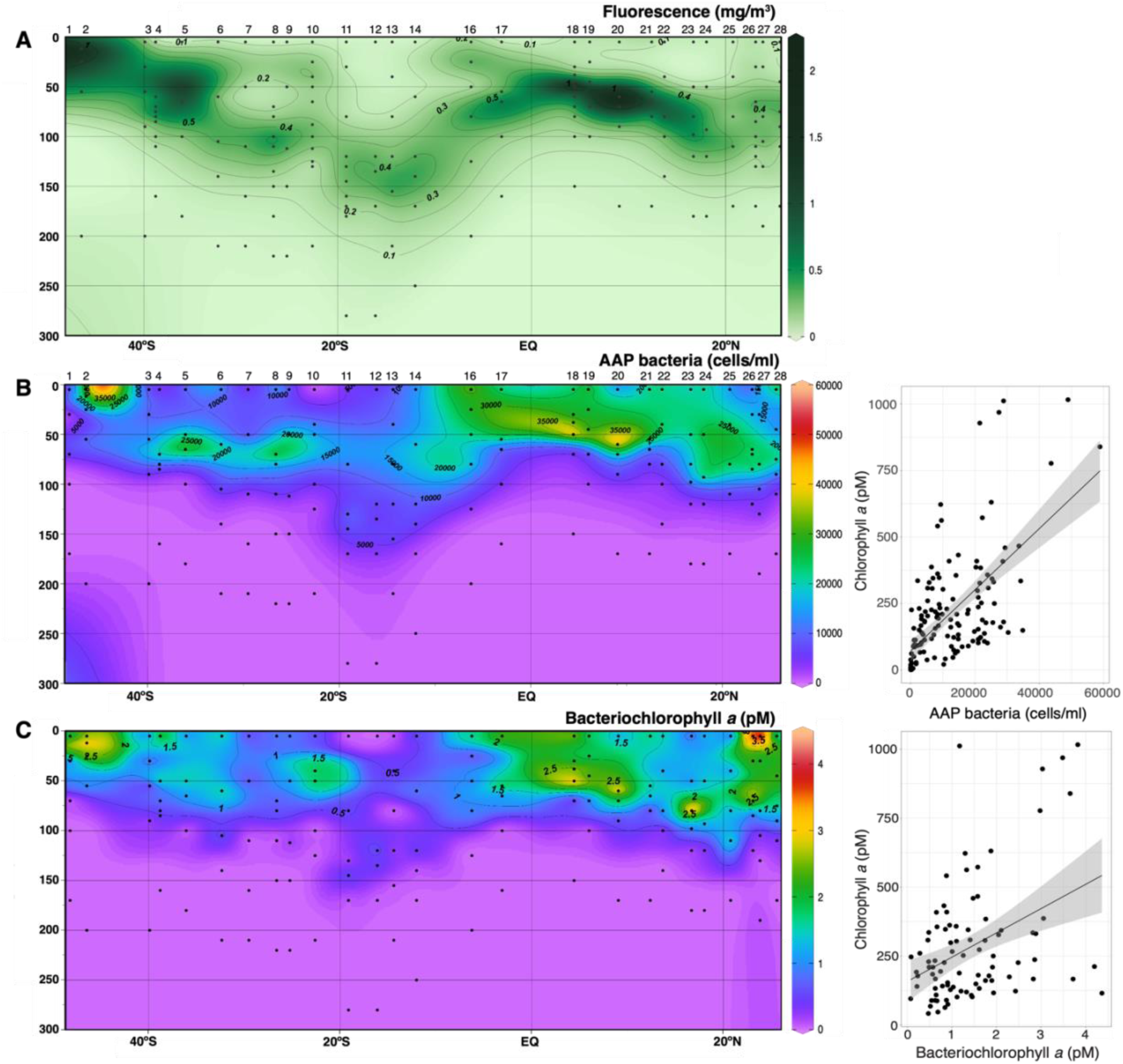
Pigment concentration and abundance distribution of aerobic anoxygenic phototrophic (AAP) bacteria along the South and Mid Atlantic Ocean. A) Sectional distribution of chlorophyll fluorescence (mg/m^3^) as obtained from the CTD/Fluorometer data. B) Sectional distribution of AAP abundance (cells/mL) estimated by epifluorescence microscopy (left) and its correlation with the chlorophyll *a* concentration (right). C) Bacteriochlorophyll *a* concentration (pM) estimated with HPLC (left) and its correlation with the chlorophyll *a* concentration (right). Station numbers are indicated above panels A and B. Black dots in the left panels indicate the depths of sampling.

### 2. Bacterioplankton abundance variation across a vertical and horizontal gradient

Bacterioplankton (i.e. bacteria and archaea) abundances obtained from the flow cytometry analyses varied between 1.1·10^5^ and 2.2·10^6^ cells/mL (mean ± standard deviation, s.d., 6.1·10^5^ cells/mL ± 4.1·10^5^) and were mostly concentrated between 0 and 75 m depth, with a very similar distribution of high nucleic acid-containing (HNA) and low nucleic acid-containing (LNA) prokaryotes in all samples (Figure S2). A high proportion of the bacterioplankton cells were *Prochlorococcus* (∼47%), which dominated in almost all stations and depths with values between 10^5^ and 10^6^ cells/mL. *Synechococcus* oscillated between 10^2^ and 10^4^ cells/mL and their abundance increased towards the boundary areas of the oligotrophic gyre (Figure S2B). Plastidic picoeukaryotes were more abundant at the DCM, in eutrophic areas, with densities up to 10^4^ cells/mL (mean ± s.d., 1.6·10^3^ cells/mL ± 2.7·10^3^). Altogether, the autotrophic picoplankton —*Synechococcus*, *Prochlorococcus* and picoeukaryotes— ranged between 6·10^4^ and 1.2·10^6^ cells/mL (mean ± s.d., 3.05·10^5^ cells/mL ± 2·10^5^, Figure S2B). The AAP abundance ranged between undetectable values below the DCM (in 7% of samples) and 5.8·10^4^ cells/mL (mean ± s.d., 1.1·10^4^ ± 1·10^4^ cells/mL), surpassing the abundances of *Synechococcus* and picoeukaryotes in most samples by an order of magnitude. In some stations within the oligotrophic gyre (stations 11 and 12), with deeper DCMs, we could detect them down to 280 m depth (our deepest sampling). AAP bacteria represented between 0.05% and 8% of total bacterioplankton (mean ± s.d., 1.5 ± 1.1 %). The highest values were observed in the eutrophic southernmost stations and in the equatorial upwelling, corresponding to samples with high concentrations of Chl *a* (Pearson correlation between untransformed AAP abundances and Chl *a*, n=172, R=0.62, *p*<0.005, and between %AAP, n=172, R=0.42, *p*<0.005, Figure 2B). This correlation was especially strong and significant in samples from the DCM and below the DCM (Pearson correlations with Chl *a*, n=27, R=0.69, *p*<0.005 and n=75, R=0.82, *p*<0.005 respectively). Interestingly, in station 10, which had a two peak DCM structure, the AAP abundance was maximal coinciding with both DCMs (Figure S3). All in all, the distribution of AAP bacteria along the water column followed the Chl *a* variation, with maximal abundances generally above the DCM and in some cases, at the DCM (Figure 2B). In most stations, the maximum AAP abundances coincided with the maximum cell abundances of the autotrophic picoplankton.

### 3. Bacteriochlorophyll *a* distribution

Bchl *a* concentration in the upper 150 m of the water column varied between 0 and 4.36 pM (mean ± s.d., 1.43 pM ± 0.98). The total concentration of Chl *a* was at least two orders of magnitude higher than that of Bchl *a* and varied between 0.89 and 1016 pM (mean ± s.d., 207.6 pM ± 205.3). As shown for AAP abundances, the distribution of Bchl *a* was highly correlated with that of Chl *a* (Pearson correlation between Bchl *a* and Chl *a*, n=172, R=0.56, *p*<0.005). When we considered only the surface samples, the correlation was weaker (n=27, R=0.36, *p*=0.09) due to the high Chl *a* concentration in samples from stations 1, 2, and 3, where we detected a diatom bloom (results not shown). When these stations were removed, Bchl *a* and Chl *a* were strongly and statistically correlated in the surface (n=24, R=0.62, *p*<0.005). In almost all samples, Bchl *a* concentration peaked above or at the DCM (Figure 2), except for a few sites (stations 1, 4, 21, and 27), where Bchl *a* was highest at the surface. The proportion of Bchl *a* in total photosynthetic pigments (Bchl *a*/(Bchl *a* + Chl *a*), ranged from 0% to 3.65%, being higher in the surface and decreasing in samples from deeper DCM layers (Figure S4). The Bchl *a* cellular quota ranged between 8.9 and 809 ag/cell (mean 49.1 ag/cell). The variation of the Bchl *a* quota correlated well with the abundance of the autotrophic picoplankton, specially the abundance of picoeukaryotes and with some inorganic nutrients such as nitrate (Figure S5).

### 4. Light energy captures by AAP bacteria and phytoplankton

The concentration of Chl *a*-containing photosynthetic units (PSUs) varied between 1.78·10^9^ to 2.04·10^12^ PSU/L, exceeding by around two orders of magnitude Bchl *a*-containing PSUs (1.34·10^9^ to 7.73·10^10^ PSU/L). Chl *a*-containing PSUs significantly increased from surface towards deeper layers (Tukey test, *p* < 0.05, Figure S5), while the concentration of Bchl *a* containing PSUs was very similar in the surface, above the DCM or in the DCM (Tukey test, *p* = 0.665), and only significantly decreased below the DCM (Figure S6), where the pigment concentration was very low. The cellular light energy captured by AAP cells ranged between 3.16·10^-12^ and 1.56·10^-9^ J/cell·day (mean 8.01·10^-11^ J/cell·day), about an order of magnitude lower than that captured by autotrophic picoplankton (mean 3.51·10^-10^ J/cell·day) (Figure 3A). The cellular light energy captured by autotrophic picoplankton cells increased from the surface to the DCM, consistent with the pattern of increasing concentration of Chl *a*-containing PSUs. On the contrary, the energy captured by AAP cells significantly decreased with depth (Figure 3, Tukey test, *p* < 0.005), since while light intensity declined, the concentration of Bchl *a*-containing PSU per cell did not vary across the different epipelagic layers, and was only significantly lower below the DCM. The divergent pattern of energy captured by both pigments along depth is evident when we represent the ratio of cellular daily energy captured by Bchl *a* and Chl *a* in each layer of the epipelagic (Figure 4). AAP bacteria capture the maximum cellular light energy in the surface, and their contribution rapidly decreases towards deeper waters. At all depths, the values of light energy captured by AAP bacteria exceeds the costs of constructing the proteins and pigments structuring their PSUs (calculated by Kirchman and Hanson (2013) to be ca. 8.29·10^-15^ J per PSU), with a mean net benefit of 1.47·10^-10^ J/cell·day. Finally, we estimated the depth-integrated energy potentially captured by AAP bacteria and by autotrophic picoplankton in the water column (Figure 4B). For both pigments, the concentration of PSU per square meter was higher at more productive regions and lower in the oligotrophic gyres. In contrast to the differences between AAP bacteria and picophytoplankton along the vertical gradient, both groups displayed the same horizonal pattern of estimated energy captured, with lowest values in the South Atlantic oligotrophic gyre that increased towards more eutrophic sites (Figure 4B, Pearson correlation of depth-integrated energy captured, n=27, R=0.49, p<0.05).

**Figure 3.**
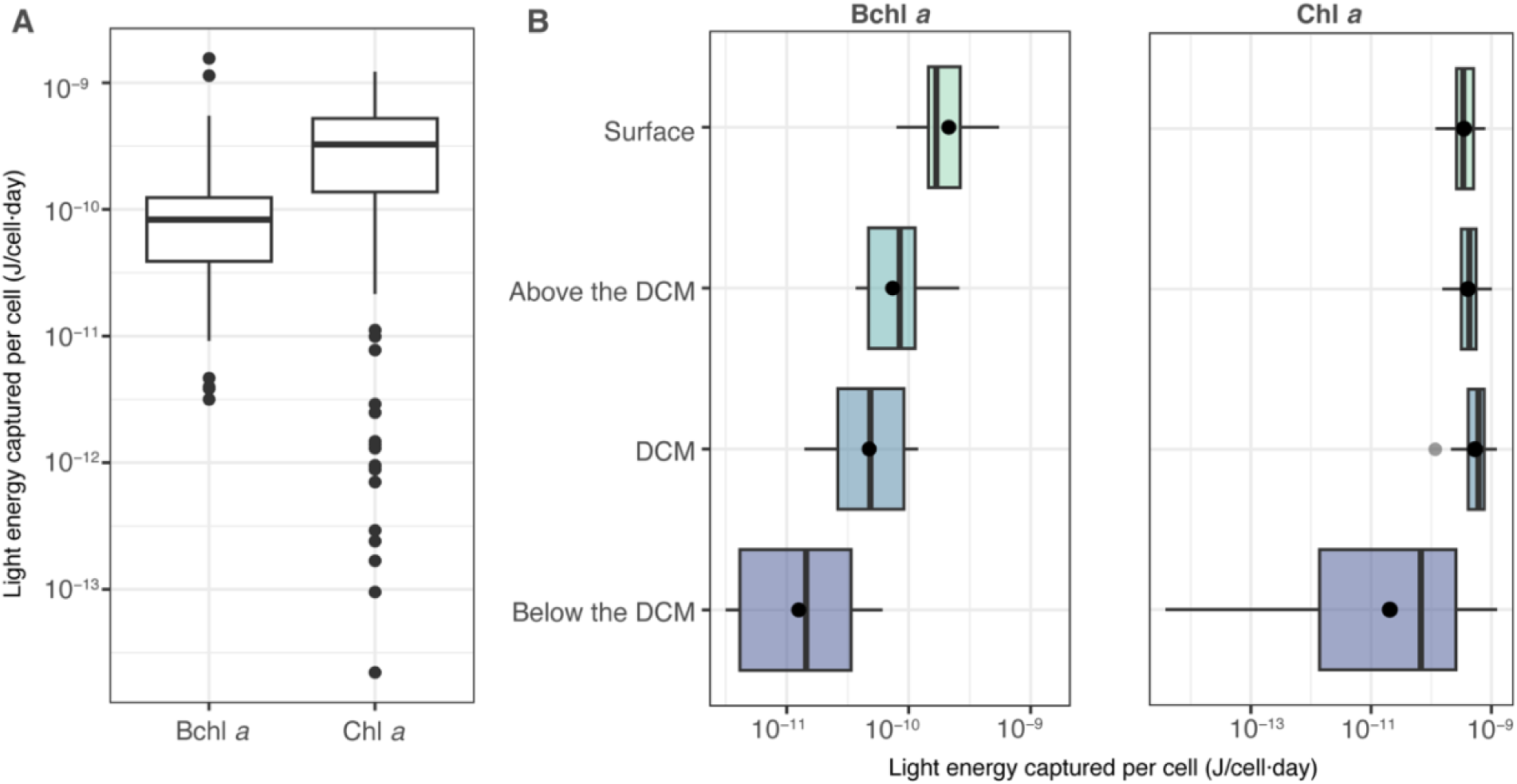
Cellular daily energy captured by bacteriochlorophyll *a* (Bchla *a*) and chlorophyll *a* (Chl *a*) along the South and Mid Atlantic Ocean. A) Light energy captured per cell and day by bacteriochlorophyll *a* and chlorophyll *a*. B) Pattern of light energy captured per cell and day along the epipelagic layers, for each pigment. The dot within the boxplots is the mean, while the thick line represents the median. Outliers are represented with dots outside the boxes.

**Figure 4.**
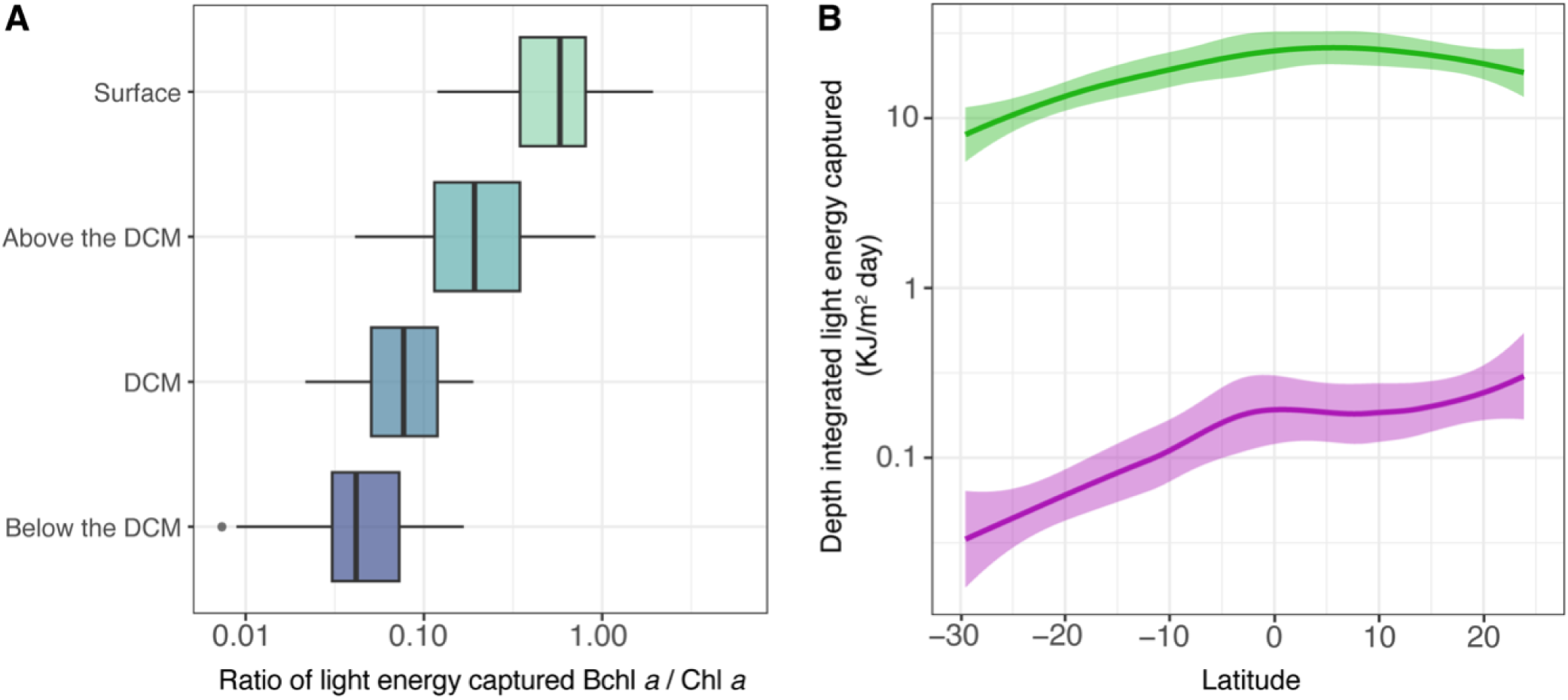
Vertical and horizonal cellular daily energy captured. A) Ratio of the cellular daily energy captured by Bchl *a* and by Chl *a* at each layer within the epipelagic and B) depth integrated light energy captured by phytoplankton (green line), and AAP bacteria (purple line) along the Atlantic Ocean transect.

## Discussion

Aerobic anoxygenic phototrophic (AAP) bacteria are ubiquitous in the sunlit zone of the ocean, where they play an important ecological role in the carbon cycle despite their relatively low abundances (Kolber et al. 2000, Kolber et al. 2001, Koblížek 2015). Although their distribution in the surface waters of different oceanic regimes has been reviewed, little information is known about their vertical distribution patterns along the photic zone. Our sampling took place in the South and Mid Atlantic Ocean, in the South hemisphere fall and Northern hemisphere spring, following a transect of contrasting productivity and nutrient concentrations. The areas with higher productivity corresponded to either the southernmost stations, influenced by the proximity to the Southern Ocean and where frequent diatom blooms are reported (Malviya et al. 2016), and the stations along the equatorial upwelling. Samples from the oligotrophic gyre could be distinguished by their low surface Chl *a* values (Figure 2A), but *Synechococcus* abundances also served as a good biological indicator of the gyre boundaries (Hartmann et al. 2012), as their concentrations decreased around an order of magnitude in samples within the oligotrophic gyre. *Prochlorococcus*, the most abundant cyanobacteria in the South Atlantic Ocean (Zubkov et al. 2000; Carvalho et al. 2020), numerically dominated all stations and samples. These cyanobacteria were the main components of the picophytoplankton, and together with picoeukaryotes, their abundances were in line with previous data estimated in this region (Visintini et al. 2021). Picoeukaryotes contribute significantly to the total C biomass in the Atlantic Ocean, as seen before by the Atlantic Meridional Transects (Tarran et al., 2006). In our samples, these three groups —*Synechococcus*, *Prochlorococcus* and picoeukaryotes—, were the main components of the phytoplankton in most of the transect, with the exception of stations 1, 2, 3 and 4, where diatoms were also an important fraction of the total phytoplankton. All in all, the sampled transect showed a high variability in terms of trophic conditions which was reflected in the DCM structures along the water column, with some DCM being more shallow and intense in terms of Chl *a* concentrations while others were located below the 100 m depth.

Along these variable vertical profiles, the abundance of AAP bacteria followed the Chl *a* distribution, peaking above or at the DCM level, and in most samples coinciding with the autotrophic picoplankton maximum abundance, confirming the outlined hypothesis that AAP bacteria would be connected to the DCM structure. Across the Atlantic Ocean, we observed the highest abundances in the eutrophic southernmost stations and in the equatorial upwelling, where the input of nutrients from deeper layers supports the productivity of phytoplankton communities. Likewise, the distribution of both Bchl *a* and Chl *a* was very similar and these pigments occupied similar areas along the water column. The concentration of Bchl *a* in our sampling was comparable to that in other marine regions like the Pacific Ocean (Kolber et al. 2001; Goericke 2002; Lami et al. 2007, Jiao et al. 2010), the Atlantic Ocean (Cottrell et al. 2006; Sieracki et al. 2006), the Baltic Sea (Koblížek et al. 2005) and the Mediterranean Sea (Lami et al. 2009; Hojerová et al. 2011; Ferrera et al. 2014; Gómez-Consarnau et al. 2019). It has been extensively documented that AAP bacteria are more abundant in more productive areas (reviewed in Koblížek, 2015), and the overlap between AAP cells and Chl *a* has been reported before in areas from the Atlantic and Pacific Oceans (Kolber et al. 2001; Schwalbach and Fuhrman 2005; Cottrell et al. 2006; Mašín et al. 2006, Jiao et al., 2007, Jiao et al., 2010), or the Mediterranean Sea (Hojerová et al. 2011), yet information along vertical profiles is scarce and mainly focused on the Northern hemisphere (for example, Sieracki et al., 2006, Mašín et al., 2006, Lami et al., 2009). The fact that Bchl *a* concentration mirrors the distribution of Chl *a* throughout depth and along the latitudinal axis, could indicate that the photoheterotrophic activity of AAP bacteria might be favoured or controlled by the same mechanisms that control phytoplankton, but it could also indicate the existence of strong biotic interactions between these two microbial guilds. In fact, co-occurrence networks have shown interactions of some AAP bacteria with phototrophic eukaryotes (Auladell et al., 2019), and various AAP species have been isolated from phytoplankton, particularly from dinoflagellates (Biebl et al., 2005, Yang et al., 2018). AAP cells with higher Bchl *a* content were associated to the most productive areas, with higher abundances of picophytoplankton or nitrate, while the light intensity across sites did not have any effect on the Bchl *a* quota. Similarly, the number of Bchl *a*-containing PSUs per cell did not vary along the surface, above the DCM and at the DCM, suggesting that no photoacclimatation processes were operating, or at least we could not detect them. In this sense, although a similar regulatory circuit based on the *ppsR* (a transcriptional regulator of photosystems) has been described in several subclasses of Alpha- and Gammaproteobacteria (Liotenberg et al. 2008; Tomasch et al. 2011; Zheng et al. 2011), different AAP strains show variable adaptations to changes in light intensity. For example, *Roseobacter litoralis* decreased three times the number of Bchl *a* molecules in the PSU following increasing irradiance (Selyanin et al. 2016), whereas *Dinoroseobacter shibae* adjusted its electron transport rate (Piwosz et al. 2018). These studies are based on cultivated AAP species and little information regarding their phototrophic activity that can be applied to natural communities exists. Thus, following the approaches by other authors (Kirchman and Hanson 2013; Gómez-Consarnau et al. 2019), we assumed a constant number of molecules per PSU and a constant electron transfer rate in our calculations.

Previous approaches estimating the light energy captured by AAP bacteria in natural environments were theoretical (Kirchman and Hanson 2013), or based solely on pigment data and hypothesized AAP abundances (Gómez-Consarnau et al. 2019). While these studies have constituted a solid base for this study, we estimated the light energy captured by AAP cells and the autotrophic picoplankton, based on in situ cell abundances and pigment concentration data, and provide, for the first time, an accurate picture of the potential phototrophy of AAP bacteria in the open ocean. Our data shows that photoheterotrophy is especially strong at the surface, where the light energy captured by AAP cells is highest (Figure 4). However, this additional source of energy gained from light cannot cover the maintenance energy costs of a large bacterium, estimated as 4·10^-10^ J (Kirchman and Hanson 2013), but it probably replaces part of the oxidative phosphorylation, as seen in previous experiments (Koblížek et al. 2010; Hauruseu and Koblížek 2012), providing a fraction of the cell metabolic needs and saving substrates that can be further used for biosynthesis (Hauruseu and Koblížek 2012). All in all, the light energy captured by Bchl *a* was about an order of magnitude lower than that captured by Chl *a*, which makes sense since AAP bacteria harvest light to supplement their heterotrophic metabolism and do not rely on phototrophy for growth as phytoplankton do. It has been estimated that about 20% of their energy requirement could be satisfied by photoheterotrophy (Kolber et al., 2001), and that this reduction in the organic carbon consumption would be equivalent to 5.4% of the primary production in oceanic waters. Taken together, these results suggest that AAP activity could reduce substantially respiration in surface waters, and, in areas where they are abundant, this could have an impact on the global carbon cycle.

In summary, we show that AAP bacteria are widespread throughout the open ocean in areas of contrasting productivity, from surface down to 250 m deep. They are strongly associated with the autotrophic picoplankton, as seen by their abundance distribution and Bchl *a* quota variation. The preference of AAP bacteria for chlorophyll rich waters could be due to common variables affecting the distribution of both functional groups but could also indicate their dependence on the phytoplankton. AAP cells are larger than average bacteria (Sieracki et al. 2006) and display higher growth rates under different scenarios (Koblížek et al. 2007; Ferrera et al. 2011, 2017). To sustain such a dynamic metabolism, they must rely on large quantities of phytoplankton-derived dissolved organic matter, and therefore they would preferably develop in eutrophic areas. The light harvesting ability provides AAP bacteria with an additional source of energy that can be used to complement their energetic needs, especially in the surface ocean.

Overall, this is the first study that uses both, in situ cell abundances and pigment data to estimate the potential solar energy captured by AAP bacteria in such an extensive area of the open ocean. The extent of our sampling area allowed the detection of large-scale patterns, making predictions on the potential phototrophic activity of AAP bacteria along the epipelagic, contributes to a better understanding of the energy fluxes in the ocean and, ultimately, in the global carbon cycle. Future studies analysing the interaction between AAP bacteria communities and the organic matter derived from different phytoplanktonic groups will shed light on the relationship between these functional groups, providing a better understanding of the ecologic role of AAP bacteria in the marine environment.

## Acknowledgments

We thank the people of the Laboratory of Anoxygenic Phototrophs in Třeboň, where the epifluorescence microscopy and HPLC analyses were performed. We are particularly grateful to Lucija Kanjer MSc. for her help during optimization of our HPLC protocol, Dr. Martí Gali for his help in obtaining the satellite data and Dr. Laura Gómez-Consarnau for her guidance in the bioenergetic calculations. We would also like to thank all the crew members of the *R/V Sarmiento de Gamboa*, as well as the scientific team, particularly Alba González-Vega, Eugenio Fraile-Nuez, and chief scientist Dr. Jesús M. Arrieta. This work was supported by grants ECLIPSE (PID2019-110128RB-I00/AEI/10.13039/501100011033) to IF and MICOLOR to JMG (PID2021-125469NB-C31) and OS (PID2021-125469NB-C32), funded by the Agencia Estatal de Investigación from the Spanish Ministry of Science and Innovation. Authors affiliated to the Institut de Ciències del Mar received the institutional support of the ‘Severo Ochoa Centre of Excellence’ accreditation (CEX2019-000928-S). CRG was supported by a PIF fellowship from the Universitat Autònoma de Barcelona.

## Author contribution

C.R.G. and I.F. conceived the study, I.F., J.M.M., O.S. and J.M.G. adquired funding, I.F. collected the samples and M.K. and C.R.G. analyzed the samples. C.R.G. processed the data and wrote the first draft. All authors contributed significantly to interpretation of results and review the manuscript.

## Data availability

Abundance and pigment data, and code for the bioenergetic calculations is available at: https://github.com/crgazulla/AAP_bacteria_Atlantic_Ocean

