## Supplementary Information for "Vertical distribution and light energy capture by AAP bacteria along contrasted areas of productivity in the Atlantic Ocean"

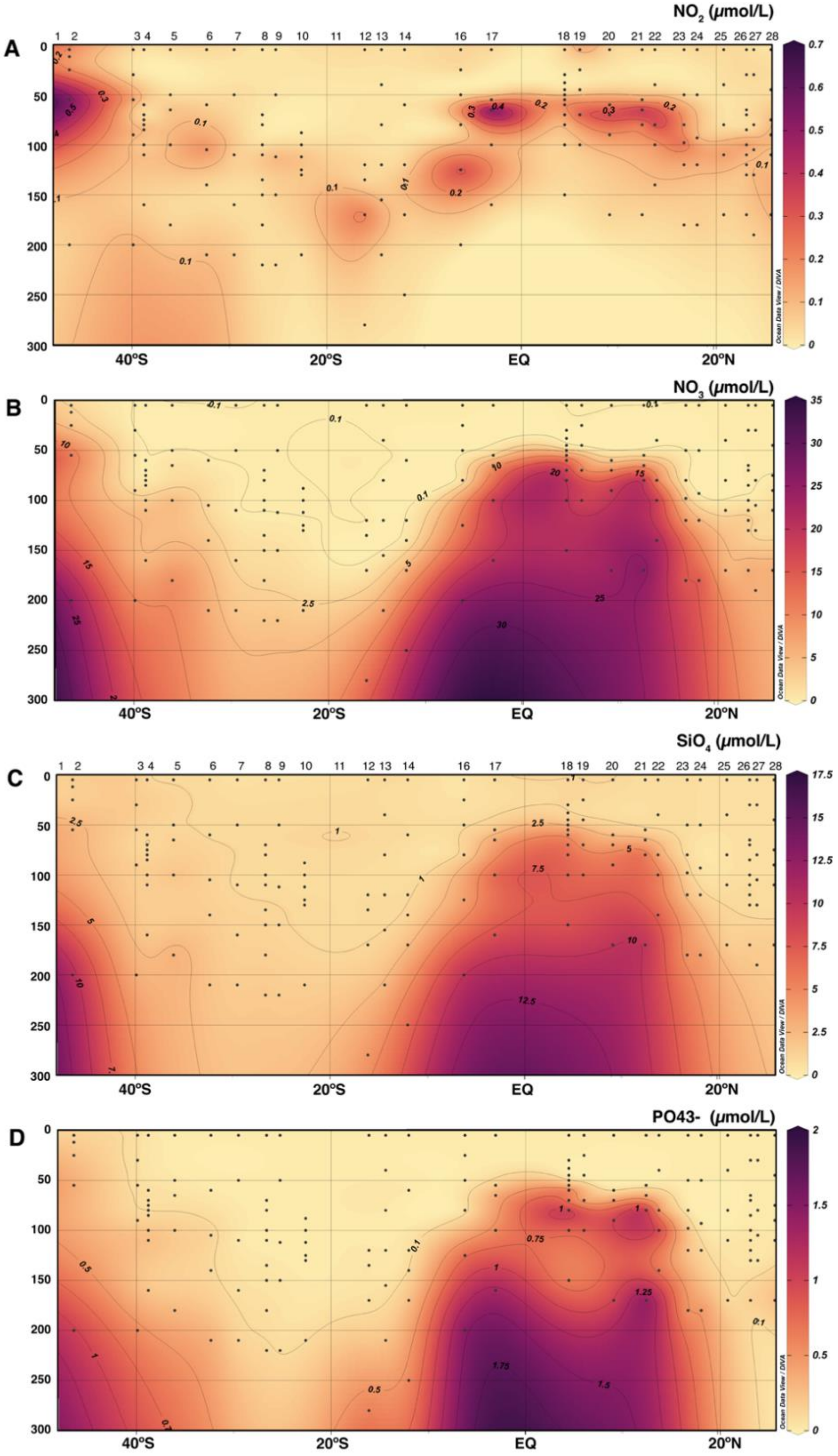


**Figure S1.** Section distribution of nitrite ($NO_2^-$), nitrate ($NO_3^-$), silicate ($SiO_4^{4-}$) and phosphate ($PO_4^{-3}$) along the South And Mid Atlantic Ocean. Black dots in all panels indicate the depths of the discrete samples.

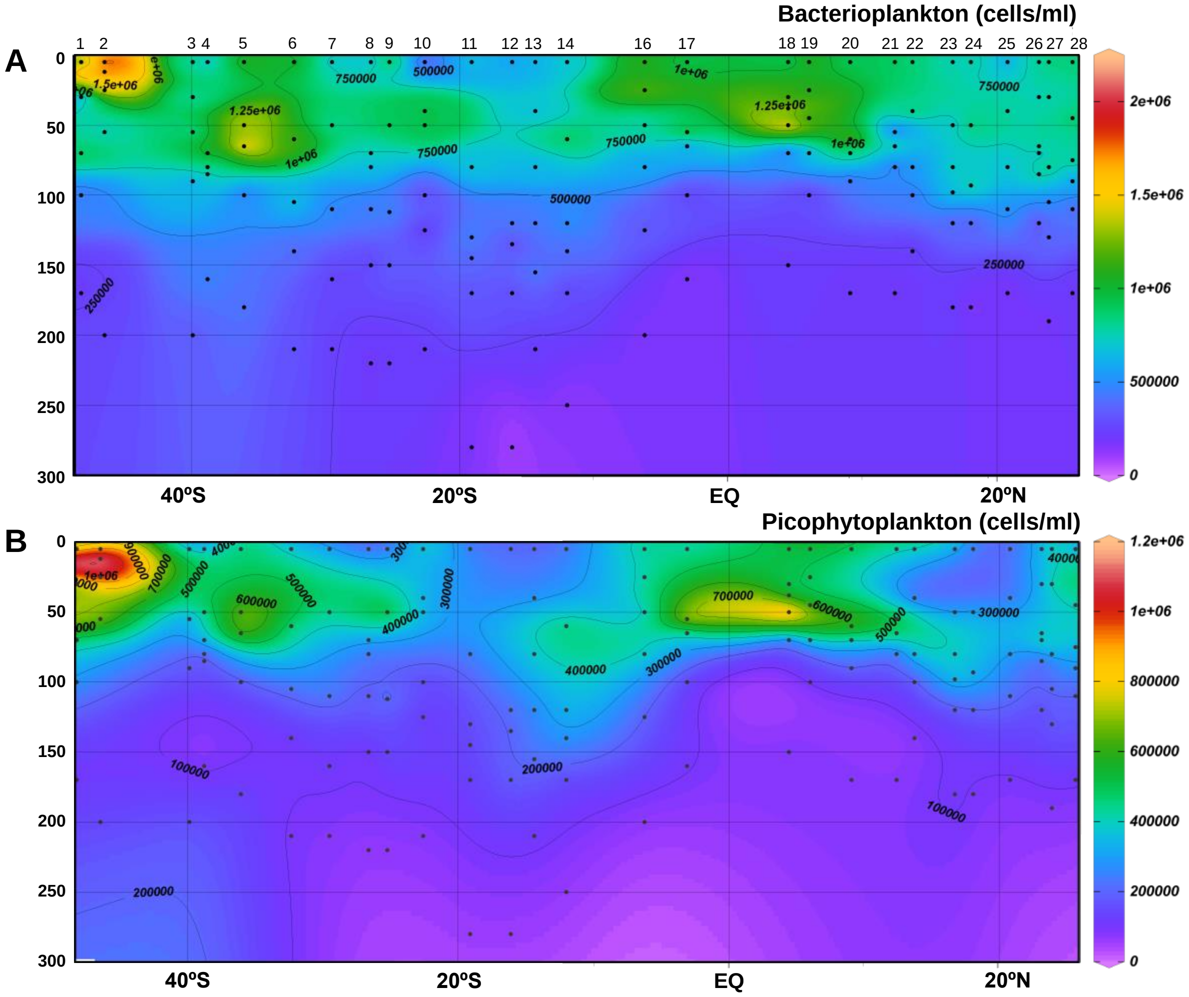


**Figure S2.** Distribution of bacterioplankton (A) and picophytoplankton (B) along the water column in the south and mid-Atlantic Ocean. Abundances of total prokaryotes are from epifluorescence microscopic analyses, after staining cells with DAPI. Picophytoplankton data is the sum of *Synechococcus*, *Prochlorococcus* and picoeukaryotes determined by flow cytometry.

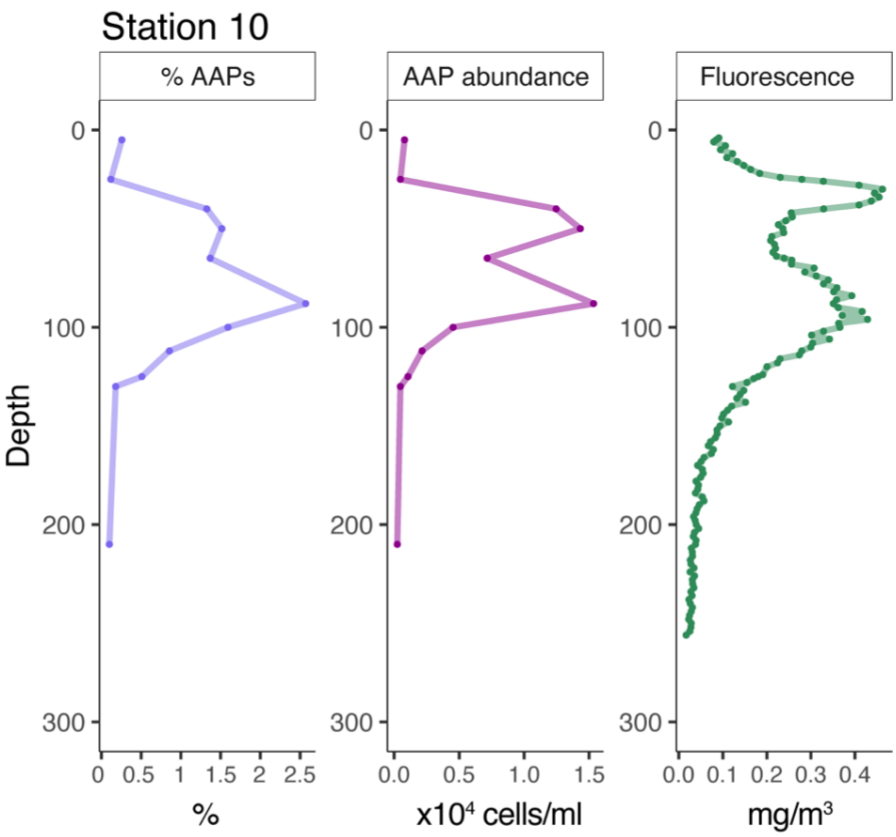


**Figure S3.** Depth profile of station 10 showing (from left to right) the percentage of AAP bacterial abundances in the bacterioplankton, the concentration of cell abundances of AAPs, and the fluorescence depicted by the CTD profiler.

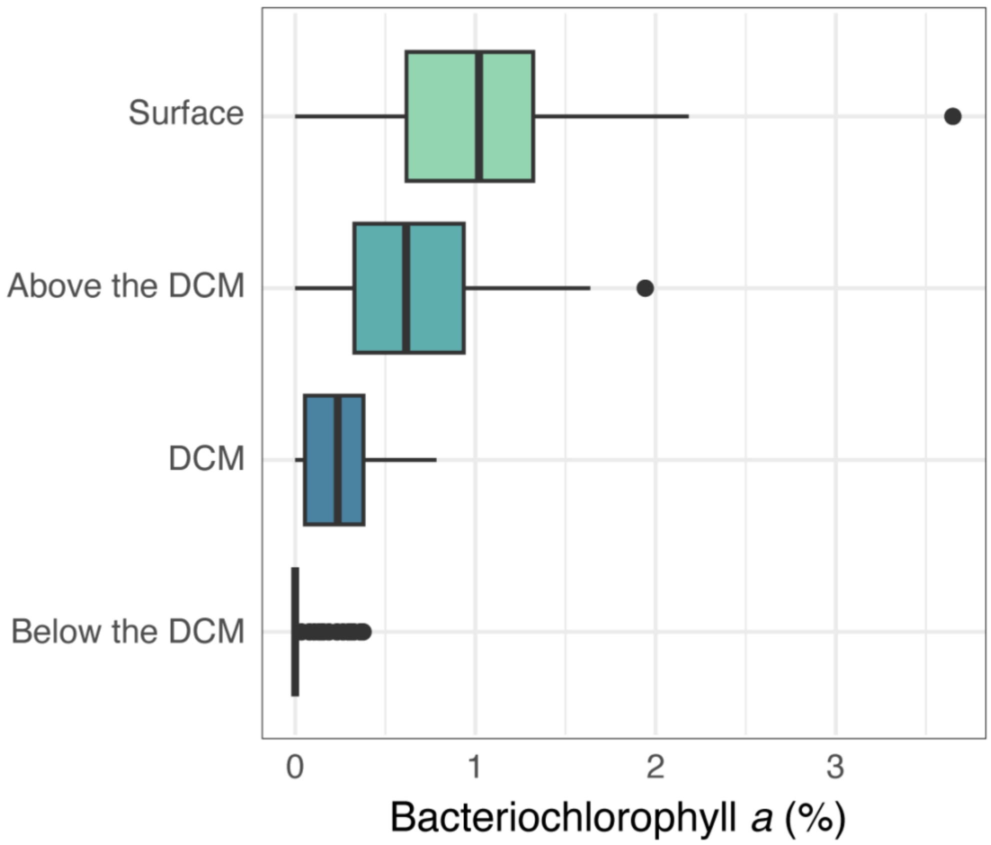


**Figure S4.** Bacteriochlorophyll *a* proportion at each layer of the epipelagic structure.

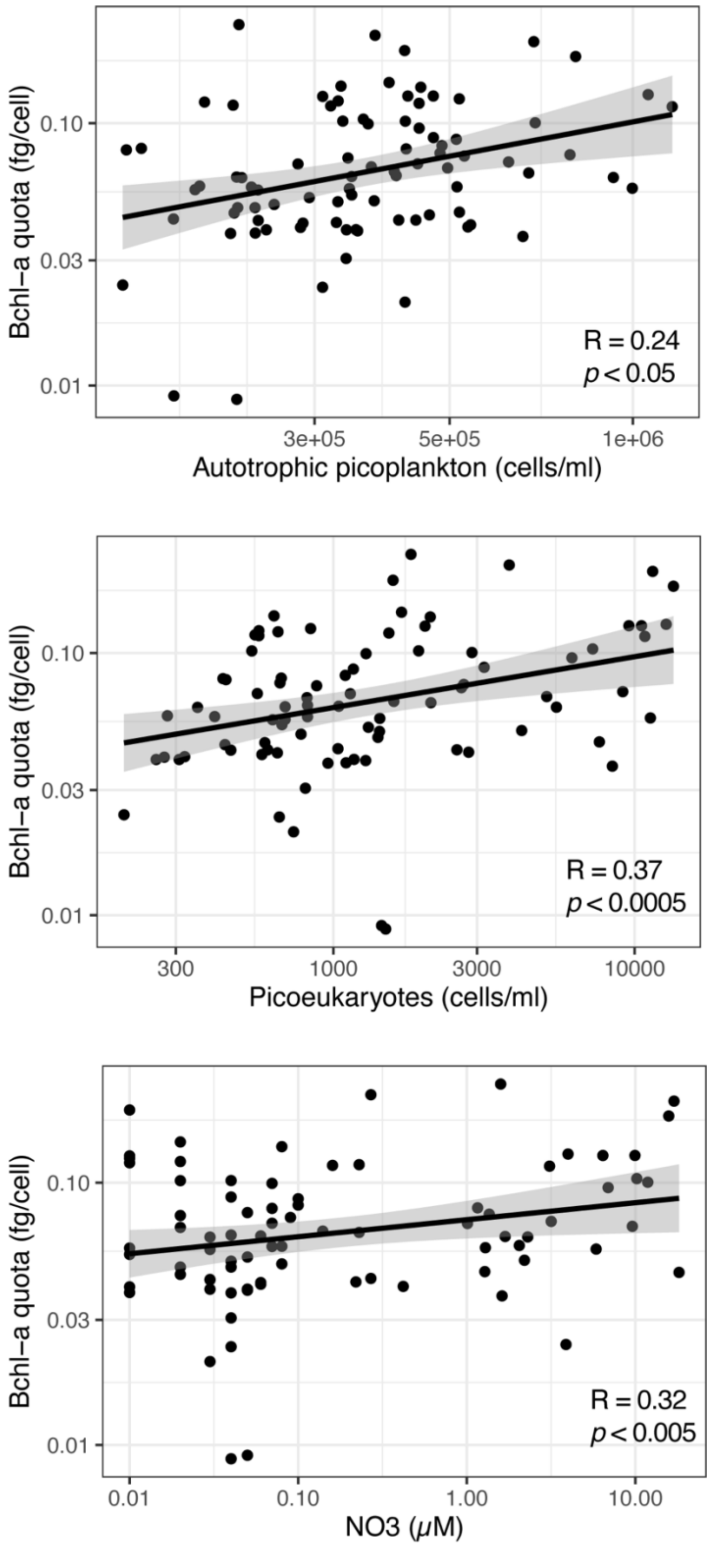


**Figure S5.** Pearson correlation between the Bchl *a* quota and (from left to right), the picophytoplankton, picoeukaryote abundance, and nitrate ($NO_3$).

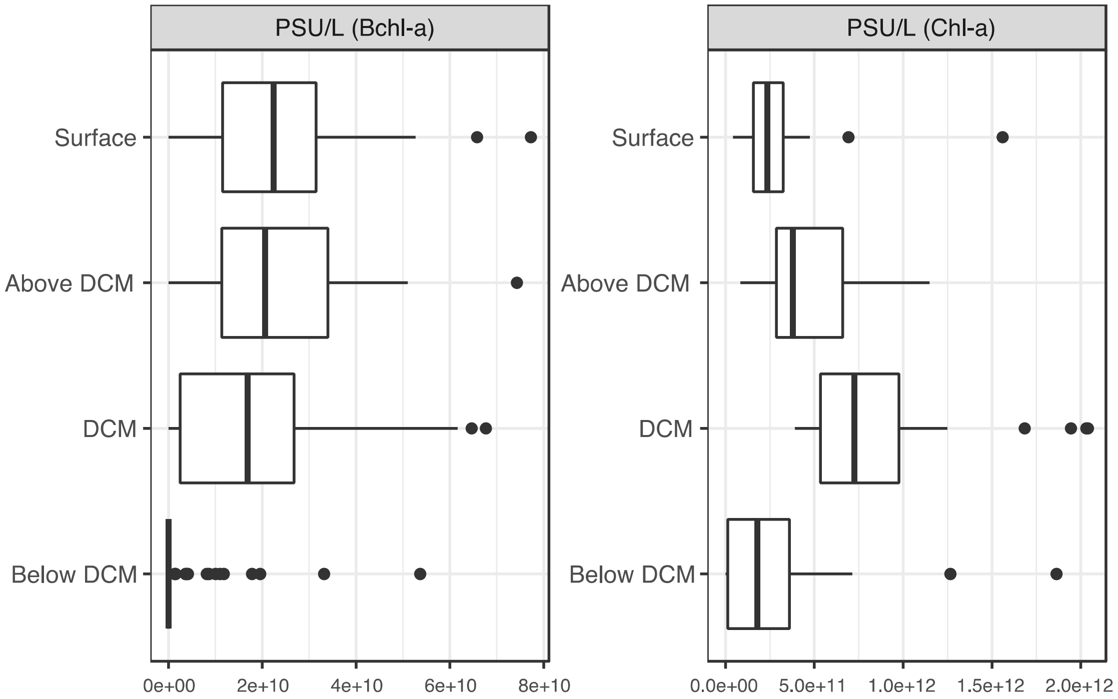


**Figure S6.** Concentration of Bchl *a* and Chl *a* based photosystem unit (PSU) along the epipelagic layers.
